# Interactive effects of herbivory and the level and fluctuations of nutrient availability on dominance of alien plants in synthetic native communities

**DOI:** 10.1101/2021.07.26.453898

**Authors:** Yanjun Li, Yingzhi Gao, Mark van Kleunen, Yanjie Liu

## Abstract

Numerous studies have highlighted the role of nutrient availability and fluctuations therein for invasion success of alien plants. Others also highlighted the role of herbivores in invasion success. However, how herbivory and the level and fluctuations in nutrient availability interact in driving alien plant invasion into resident communities remains largely unexplored.

We grew eight invasive alien species as target species in pot□mesocosms with five different synthetic native communities in a three-factorial design with two levels of nutrient availability (low *vs* high), two levels of nutrient fluctuations (constant *vs* pulsed) and two levels of herbivory (with *vs* without).

The relative biomass production of the alien target plants decreased in response to an increase in nutrient availability, and increased in response to the presence of herbivores. Furthermore, herbivory could interact with changes in nutrient availability and nutrient fluctuations to affect the dominance of the alien target species (a marginally significant interaction; 90% CIs: [0.125, 2.712]).

Our multispecies experiment indicates that herbivory could mediate the interactive effect of nutrient enrichment and variability in nutrient supply on invasion of alien plants into native communities. Therefore, we recommend that studies testing the fluctuating resources hypothesis should also consider interactive effect of other trophic levels.

## Introduction

Invasion by alien plants could reduce native biodiversity, influence ecosystem functions, and degrade ecosystem services [1]. Due to the rapid globalization, the increase in the number of naturalized alien plant species does not show any sign of saturation, and it was recently predicted that their numbers may increase on average by 18% from 2005 to 2050 [2–4]. Therefore, a major research objective in the field of ecology is to identify the mechanisms that underlie alien plant invasion [5–9].

It has frequently been suggested that an increase in soil-nutrient availability is one of the most important drivers of alien plant invasion [8, 10, 11]. As successful alien plant species are often introduced from anthropogenic and more nutrient-rich environments, they might be more likely to be adapted to high-nutrient environments [12, 13]. Indeed, both a recent meta-analysis [14] and a globally replicated study of 64 grasslands [15] showed that successful alien plants respond more strongly to nitrogen enrichment than native plants do. So far, empirical studies that tested how soil-nutrient changes affect alien plant invasion mainly focused on changes in the mean nutrient level [16–22]. However, due to increasing occurrences of extreme events (i.e., droughts, floodings, heat waves, fires), soil-nutrient changes also entail changes in their variability, and this may affect plant invasions [20, 23]. Therefore, it is important to test how changes in soil-nutrient levels, as well as fluctuations in nutrient availability over time, drive alien plant invasion in resident communities.

The fluctuating resource hypothesis proposes that temporal fluctuations in nutrient supply could promote alien plant invasion in resident communities [10]. However, empirical studies testing the hypothesis found mixed results. For example, Parepa et al. found that a pulsed nutrient supply, compared to a constant nutrient supply, increased the dominance of the invasive plants *Fallopia japonica* and *F. × bohemica* in experimental plant communities [23]. In contrast, Liu et al. showed that a pulsed nutrient supply decreased the dominance of invasive alien plants [24]. Thus, more studies are needed to test the hypothesis, and why the results might vary. The studies that tested the effect of temporal changes in nutrient availability did so under overall nutrient-rich conditions. Under more nutrient-limiting conditions, however, the effect of temporal fluctuations may be even stronger. In other words, mean nutrient availability may interact with temporal fluctuations in nutrient availability to affect alien plant invasion into resident communities. However, very few studies have tested whether this expectation holds [25].

Although the fluctuating resource hypothesis has become a key theory in invasion ecology, previous tests only used study systems consisting of a single trophic level (i.e., only considered plant-plant interactions). Plant growth, however, can be strongly regulated by other trophic levels, such as herbivores. This might be relevant for the fluctuating resource hypothesis as alien species are likely to be released from most of their native enemies, and thus should suffer less herbivory than native species in their introduced regions [26–28]. Following this logic, the presence of herbivores, just like increases in resource availability and fluctuations therein [20, 23, 29, 30], could promote alien plant invasion in resident communities. Moreover, as herbivore effects on plants are often regulated by soil-nutrient availability, because plants growing in relatively high-nutrient conditions might be better able to compensate or tolerate herbivory [31–34], the effects of herbivores might interact with the effects of nutrients. Therefore, it is reasonable to assume that the presence of herbivores might amplify the positive effect of increases in resource availability and fluctuations on alien plant invasion.

To test the individual effects of nutrient availability, nutrient fluctuations, herbivory, and their interactions on alien plant invasion into resident communities, we grew eight invasive alien species as target species in pot mesocosms with five different synthetic native communities, each consisting of three grassland species. Then, we exposed the plants to eight combinations of two nutrient availability (low *vs* high), two nutrient-fluctuation (constant *vs* pulsed) and two herbivory (with *vs* without) treatments. By comparing the absolute aboveground biomass production of the alien target species as well as their biomass production relative to the biomass production of the native competitors, we addressed the following specific questions: 1) Do nutrient availability, nutrient fluctuations and the presence of herbivores promote the absolute and relative biomass of alien plants? 2) Does the effect of nutrient fluctuations on absolute and relative biomass of alien plants depend on the overall nutrient availability level? 3) Does the presence of herbivores interact with increases in nutrient availability and fluctuations therein to affect the absolute and relative biomass of alien plants?

## Material and methods

### Study species

To investigate the individual and interactive effects of nutrient availability, nutrient fluctuations and the presence of herbivores on alien plant invasion into resident communities, we chose eight invasive alien species as targets, and 15 native species as native community members from the herbaceous flora of China (Table S1). We classified these species as invasive alien or native to China based on information in the book “The Checklist of the Alien Invasive Plants in China” [35] and the Flora of China database (www.efloras.org). To cover a wide taxonomic breadth, we selected the eight alien target species from seven genera of three families. As plants with different life histories (i.e., annuals or perennials) may respond differently to nutrient availability and fluctuations [20, 36], we assured that both the alien targets and native community members included annuals and perennials (Table S1). Seeds of these species were collected in natural populations in grasslands in China or ordered from a commercial seed company (see Table S1).

To impose a herbivory treatment, we selected two aboveground insect herbivores. As natural systems usually include both generalist and specialist insect herbivores, we chose the generalist grasshopper *Stenocatantops splendens* and the specialist grasshopper *Locusta migratoria* (i.e., grass-feeder) as the shoot herbivores. The grasshoppers were acquired from a commercial insect company (Cangzhou Grasshoppers Breeding Center, China). As both grasshoppers and all plant species occur mainly in grasslands, and according to the GBIF database (www.gbif.org), overlap in their distributions, they are very likely to co-occur in nature.

### Pre-cultivation and experimental setup

We conducted the experiment at the Northeast Institute of Geography and Agroecology, Chinese Academy of Sciences (43°5’49”N, 125°24’40”E). From 7 May to 26 June 2020, we sowed each of the invasive alien and native species separately into plastic circular trays (diameter = 25.5 cm, height = 4 cm) filled with potting soil (Pindstrup Plus, Pindstrup Mosebrug A/S, 103 Denmark; pH 6; 120 mg/L N; 12 mg/L P; 400 mg/L K; 28 mg/L Mg; 0.4 mg/L B; 2 mg/L Mo; 1042 mg/L Cu; 3 mg/L Mn; 0.9 mg/L Zn; 8 mg/L Fe). Because the time required for germination varies among the species, we sowed them at different times (Table S1) to ensure that at transplanting the seedlings were in a similar developmental stage. All trays with seeds were kept in a greenhouse (temperature: 18-27°C; natural lighting with an intensity of ~75% of the light outdoors; and ~68% relative humidity).

On 10 July 2020, we selected similar-sized seedlings from each of the eight invasive alien and 15 native species, and transplanted them into 2.5□L circular plastic pots (top diameter: 18.5 cm, bottom diameter: 12.5 cm, height: 15 cm) filled with a 1:1 mixture of sand and fine vermiculite. We transplanted one seedling of an alien target species in the center of each pot. For each of the eight alien target species, we transplanted a total of 40 seedlings into 40 pots (i.e., one individual per pot), resulting in total of 320 pots. Immediately after transplanting the alien target species, we equally distributed the 40 pots of each alien target over five different native communities (i.e., eight pots per native community). To create the five different native grassland communities, we randomly assigned the 15 native species into five groups of three species (Table S1). We planted two seedlings of each native community member, so that each pot included six individuals of native species equally spaced in a circle (diameter = 10 cm) around the alien target seedling. The two individuals of the same species were planted at opposite positions of the circle (Fig. 1a). After transplanting, we randomly assigned all pots to two cages (3.5 × 4.5 × 2.5 m) located outside of the pre-cultivation greenhouse. The cages were covered with transparent plastic roofs and white nylon net (mesh size: 0.25 × 0.25 mm) all around. Half of the pots of each combination of an alien target species and a specific native community were assigned to one cage, and the remaining ones to the other cage (Fig. 1a). In other words, each cage included 160 pots in total (8 alien target species × 5 native communities × 4 nutrient-supply treatments [2 nutrient availability × 2 nutrient fluctuation treatments]). To avoid the loss of water and nutrient solution, we put a plastic tray under each pot. We re-randomized the positions of the pots in each cage after five weeks.

**Figure 1.**
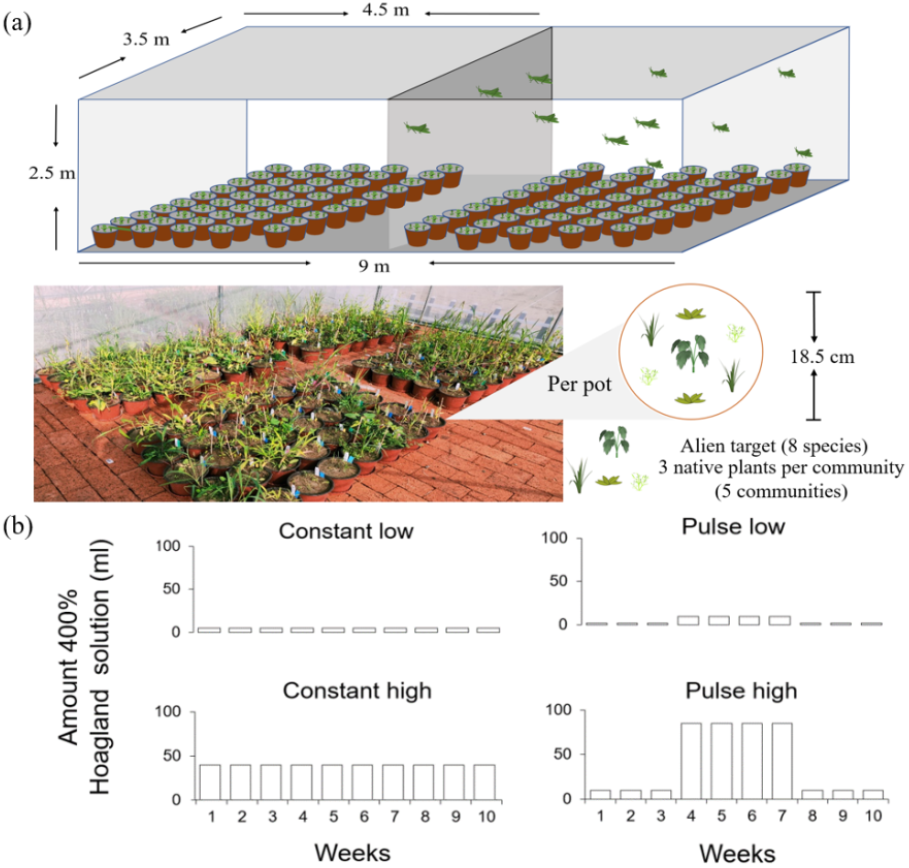
Graphical illustration of the experimental design. Overview of the herbivory treatment, the herbivory-treatment cage during the experiment, and the pot design (a); the amount of nutrient solution supplied each week during the ten weeks of the experiment (b). The constant and pulsed nutrient supply under low- or high-nutrient availability received the same total amount of nutrients during the ten weeks.

On 14 July 2020 (i.e., one week after transplanting), we started to apply the nutrient treatments at weekly intervals for a total of 10 weeks (Fig. 1b). We applied two nutrient availability treatments (low *vs* high) crossed with two nutrient-fluctuation treatments (constant *vs* pulsed), using a 400%□strength Hoagland solution (Methods S1). During the experiment, we added a total of 50 and 400 ml of the Hoagland solution to the low and high-nutrient availability pots, respectively. Although it is difficult to compare the absolute nutrient levels in a pot experiment to those found in a natural system, Liu et al. previously showed that both the low- and high-nutrient availabilities are limiting plant growth, as is usually the case in nature [24]. Within each nutrient-availability treatment, we created two different nutrient-supply patterns, a constant nutrient supply and a pulsed nutrient supply (Fig. 1b). We supplied 5 ml of the nutrient solution for the constant low-nutrient supply each week, and 40 ml of the same nutrient solution for the constant high-nutrient supply. The pulsed treatment at low-nutrient availability consisted of three weeks of 2 ml per week, followed by four weeks of 9.5 ml per week, and again three weeks of 2 ml per week (Fig. 1b). The pulsed treatment at high-nutrient availability consisted of three weeks of 10 ml a week, followed by four weeks of 85 ml, and again three weeks of 10 ml (Fig. 1b). To avoid differences in water supply among the four treatments, we added extra water to the amount of nutrient solution in each treatment to ensure that each pot received a total of 85 ml of water per nutrient application. In each cage, there were five replicates per alien target species for each of the four nutrient-supply treatments (i.e., one replicate for each of the five native communities). We watered all plants regularly by filling the dish under each pot to avoid water limitation.

The herbivory treatment started on 14 August 2020, and ended six weeks later on 20 September 2020. We added the grasshoppers in one of the two cages, and treated the other cage as control (Fig. 1a). As the commercial company hatched these two species of grasshoppers at different times, we added *Locusta migratoria* from 14 August to 6 September 2020, and *Stenocatantops splendens* from 31 August to 6 September 2020. We checked the herbivory pressure every day to determine whether we would add more grasshoppers. During the experiment, we added *Locusta migratoria* seven times (three times for 3–4^th^ instars and four times for adults), and *Stenocatantops splendens* two times (one time for 3–4^th^ instars and the other time for adults). In total, we added 354 individuals of *Locusta migratoria*, and 450 individuals of *Stenocatantops splendens* in the herbivory-treatment cage.

### Measurements

On 25 September 2020 (i.e., eleven weeks after transplanting), we harvested the aboveground biomass of all pots. For each pot, we first harvested the alien target species and then the three native competitor species. As some alien target and native plants died and three pots had accidentally been treated with the wrong nutrient solution, we only harvested 274 instead of 320 pots (see the raw data on https://doi.org/10.5061/dryad.fj6q573vn) [37]. We did not harvest the roots of the alien and native species, because the roots of the species were intertwined, and it was impossible to separate them. All aboveground biomass samples were dried at 65°C for 72 hours and then weighed. We calculated total aboveground biomass per pot by summing the biomass of the alien target species and the three native competitors. We also calculated the biomass proportion of the alien target species in each pot as the ratio between the biomass of the alien target species and the total aboveground biomass per pot.

### Statistical analysis

To test the effects of nutrient availability (low *vs* high), nutrient fluctuations (constant *vs* pulsed), herbivory treatments (with *vs* without) and their interactions on aboveground biomass production of the alien target species, biomass production of the native communities and biomass proportion of the alien target species in each pot, we fitted Bayesian multilevel models using the function brm of the R package brms [38] in R 4.0.2 [39]. In all models, we included the following explanatory variables nutrient availability (low *vs* high), nutrient fluctuations (constant *vs* pulsed) and herbivory treatment (with *vs* without) as fixed factors. To account for phylogenetic non-independence of species belonging to the same family and for non-independence of replicates of the same species, we included identity of the alien target species nested in their family as random factors in all models. To account for variation among the five different native communities, we also included identity of the native community as random factor in all models. To relax the homogeneity of variance assumption in all models, we allowed the residual standard deviation sigma to vary by the identity of alien target species [40].

For all models, we used the default priors set by the brms package, and ran four independent chains. The number of total iterations per chain was 5000, and the number of warm-up samples was 2500. To directly test hypotheses about the main effects and interactive effects based on each coefficient’s posterior distribution, we used the sum coding, which effectively ‘centers’ the effects to the grand mean (i.e., the mean value of all data observations) [41]. To implement this in brms, we used the functions contrasts and contr.sum of the stats package in R. We considered the fixed effects nutrient availability, nutrient fluctuation and herbivory treatments, and their interactions as significant when their 95% credible interval of the posterior distribution did not overlap zero, and as marginally significant when the 90% credible interval did not overlap zero. As we had only two cages available, one with herbivores and one without herbivores, our herbivory treatment is obviously pseudoreplicated [42]. This means that the main effect of herbivory should be interpreted with care [43]. However, it still allows us to test whether the effects of the nutrient availability and nutrient fluctuation treatments differ between the cage with and without herbivores.

## Results

Averaged across the nutrient fluctuation and herbivory treatments, an increase in nutrient availability significantly increased the biomass production of alien target species (+600.7%; Table 1; Fig. 2 and Fig. S1) and the biomass production of native communities (+601.0%; Table 1; Fig. 2 and Fig. S1). The presence of herbivores, however, significantly decreased the biomass production of native communities (−39.4%; Table 1; Fig. 2 and Fig. S2). The similar pattern was found for the biomass production of alien target species (−44.3%; Table 1; Fig. 2 and Fig. S2), although it was only marginally significant (90% CIs: [−0.170, −0.010]). The negative effect of herbivory on the biomass production of both alien target species and native communities was stronger under high-nutrient availability (alien: −46.4%; native: −39.7%) than under low-nutrient availability (alien: −28.0%; native: −36.6%; significant NA × H interactions in Table 1; Fig. 2 and Fig. S3).

**Table 1.** Output of the models testing effects of nutrient availability (low *vs* high), nutrient fluctuations (constant *vs* pulsed), herbivory treatments (with *vs* without), and their interactions on aboveground biomass production of the alien target species, biomass production of the native communities and biomass proportion of the alien target species in each pot. Shown are the model estimates and standard errors as well as the lower (L) and upper (U) values of the 95% and 90% credible intervals (CI).

|  |  | Estimate | SE | L95% CI | U95% CI | L90% CI | U90% CI |
| --- | --- | --- | --- | --- | --- | --- | --- |
| **Biomass production of alien target plants** | Intercept | 1.105 | 0.786 | -0.506 | 2.634 | -0.105 | 2.279 |
|  | Nutrient availability (NA) | 0.416* | 0.058 | 0.303 | 0.532 | 0.322 | 0.515 |
|  | Nutrient fluctuation (NF) | 0.046 | 0.047 | -0.047 | 0.139 | -0.032 | 0.124 |
|  | Herbivory treatment (H) | -0.090† | 0.049 | -0.185 | 0.005 | -0.170 | -0.010 |
|  | NA×H | -0.113* | 0.048 | -0.208 | -0.018 | -0.190 | -0.035 |
|  | NF×H | -0.009 | 0.048 | -0.104 | 0.086 | -0.087 | 0.071 |
|  | NA×NF | 0.031 | 0.049 | -0.066 | 0.127 | -0.050 | 0.111 |
|  | NA×NF×H | 0.020 | 0.049 | -0.076 | 0.116 | -0.060 | 0.100 |
| **Biomass production of native communities** | Intercept | 3.529* | 1.154 | 1.114 | 5.759 | 1.609 | 5.321 |
|  | Nutrient availability (NA) | 3.252* | 0.110 | 3.033 | 3.467 | 3.069 | 3.434 |
|  | Nutrient fluctuation (NF) | 0.023 | 0.111 | -0.191 | 0.238 | -0.158 | 0.204 |
|  | Herbivory treatment (H) | -1.150* | 0.109 | -1.361 | -0.939 | -1.328 | -0.973 |
|  | NA×H | -0.735* | 0.110 | -0.951 | -0.520 | -0.913 | -0.554 |
|  | NF×H | -0.144 | 0.109 | -0.359 | 0.071 | -0.324 | 0.035 |
|  | NA×NF | -0.044 | 0.109 | -0.254 | 0.167 | -0.224 | 0.133 |
|  | NA×NF×H | -0.086 | 0.109 | -0.302 | 0.129 | -0.267 | 0.095 |
| **Biomass proportion of alien target plants** | Intercept | 23.129* | 11.495 | 0.112 | 46.144 | 4.487 | 41.768 |
|  | Nutrient availability (NA) | -1.890* | 0.766 | -3.387 | -0.380 | -3.127 | -0.623 |
|  | Nutrient fluctuation (NF) | 0.649 | 0.753 | -0.820 | 2.125 | -0.592 | 1.897 |
|  | Herbivory treatment (H) | 4.229* | 0.882 | 2.470 | 5.937 | 2.754 | 5.647 |
|  | NA×H | -1.064 | 0.766 | -2.554 | 0.435 | -2.321 | 0.182 |
|  | NF×H | -0.57 | 0.765 | -2.079 | 0.921 | -1.835 | 0.665 |
|  | NA×NF | 0.918 | 0.770 | -0.636 | 2.419 | -0.343 | 2.188 |
|  | NA×NF×H | 1.432† | 0.786 | -0.123 | 2.959 | 0.125 | 2.712 |

**Figure 2.**
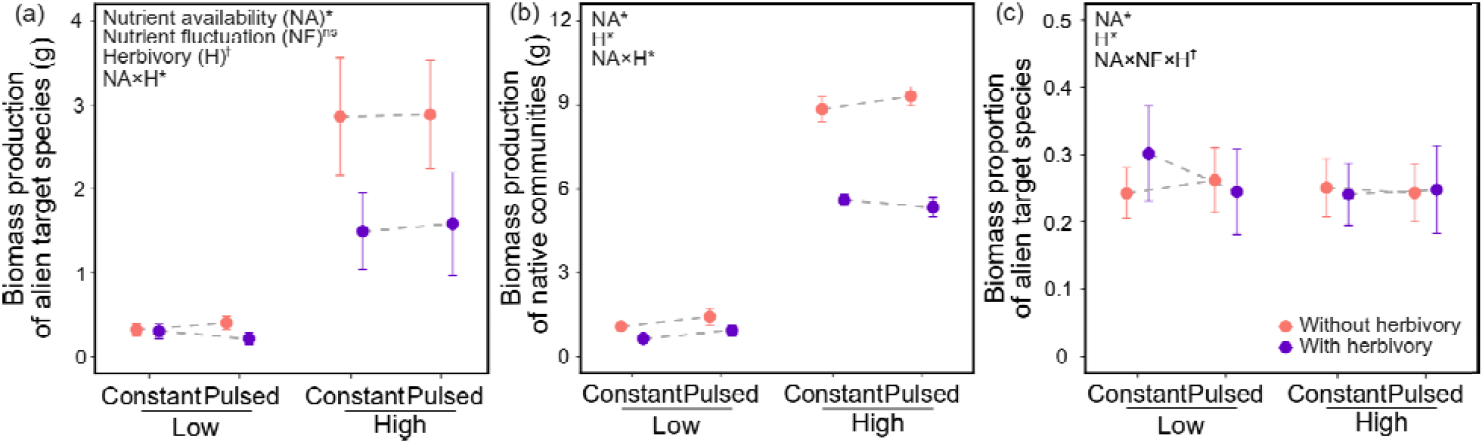
Mean values (±SE) of biomass production of alien target species (a) and native communities (b) and biomass proportion of the alien target species (c) under each combination of two nutrient availability (low *vs* high), two nutrient-fluctuation (constant *vs* pulsed) and two herbivory (with *vs* without) treatments. Parameters whose 95% credible intervals do not overlap with zero are indicted with asterisks (*), whose 90% credible intervals do not overlap with zero are indicated with daggers (†), and whose 90% credible intervals do not overlap with zero are indicated with “ns”.

We found that an increase in nutrient availability significantly decreased (−6.5%; Table 1; Fig. 2 and Fig. S1), whereas the presence of herbivores significantly increased the relative biomass production of the alien target species in the native communities (+3.6%; Table 1; Fig. 2 and Fig. S2). Additionally, under low-nutrient availability, the pulsed nutrient supply tended to increase the relative biomass production of the alien target plants in the absence of herbivores (+7.9%), whereas the reverse was true for plants in the presence of herbivores (−18.8%; Fig. 2). Under low-nutrient availability, however, pulsed nutrient supply and the presence of herbivores did not affect the relative biomass production of the alien target species (Fig. 2). The dependency of the effect of the nutrient fluctuation treatment on the levels of the other treatments was indicated by a marginally significant interaction between herbivory, nutrient availability and fluctuation therein (90% CIs: [0.125, 2.712]) in Table 1.

## Discussion

Our multispecies experiment showed that an increase of nutrient availability promoted the biomass production for both the alien target species and native communities. However, herbivory suppressed the biomass production of plants, in particular under high-nutrient availability. We found that increases in nutrient availability suppressed, whereas herbivory promoted the dominance of the alien target species in native resident communities. Interestingly, we also found tentative evidence that herbivory could interact with changes in nutrient availability and nutrient fluctuations to affect the dominance of the alien target species. In other words, herbivory could mediate the interactive effect of nutrient enrichment and fluctuations in nutrient supply on alien plant invasion into native communities.

Although each pot had one invasive alien and six native plants, the invasive plant accounted for about one quarter of the aboveground biomass in each pot (Fig. 2c). This suggests that the invasive alien species were more dominant than the native species. We also found that this dominance decreased with an increase of the average nutrient availability. This finding does not support the idea that increased nutrient availability could promote alien plant invasion in resident communities, although many theoretical [10, 44] and empirical studies [14, 15, 20, 23, 25] found evidence for this. A potenial reason could be that in our study the native species themselves are also quite common, and it are usually the rare species that take less advantage of increased nutrient availability [45].

Previous empirical studies testing the nutrient-fluctuation effect on alien plant invasion found mixed results [20, 23–25, 46, 47]. We hypothesized that this might be because the effect of temporal fluctuations may be even stronger under more nutrient-limiting conditions than under less nutrient-limiting conditions. However, we found no significant interactive effect of nutrient availability and fluctuations therein on the aboveground biomass and dominance of alien target plants in native communities. This is in line with the results of a recent case study by Gao et al. [25]. It is worth noting though that in the absence of herbivory, the absolute and relative aboveground biomass of the invasive alien plants tended to be higher in the low, pulsed nutrient treatment than in the low, constant treatment, while the reverse was true in the presence of herbivory.

Not surprisingly, herbivory decreased the biomass production of plants (also see the total biomass production per pot in Fig. S4a and Fig. S4b). In particular, the biomass suppression of herbivory was stronger under high-nutrient availability than under low-nutrient availability (Fig. S4c). This seems inconsistent with previous findings that plants compensate or tolerate herbivory more when growing in high nutrient conditions [31, 32, 48–51]. However, it should be noted that in our experiment, the herbivores could choose between the plants. As the plants grown at high-nutrient availabilities might be more nutritious [52–55], and have decreased plant secondary metabolite concentrations such as tannins [56, 57], the herbivores might have preferably fed on the plants with high nutrient availabilities [53, 58–61]. On the other hand, we found that herbivory increased the dominance of the alien target species in the native resident communities. This is also not surprising, because according to the enemy release hypothesis, alien plants often have escaped from many of their herbivorous enemies, and therefore could in their introduced range outcompete resident plants [26].

When growing under low-nutrient availability and also in the absence of herbivores, the nutrient pulse promoted the dominance of alien plants, which supports the fluctuating resource hypothesis [10]. However, when growing under low-nutrient availability but in the presence of herbivores, the nutrient pulse suppressed the dominance of alien plants in pots. This may because herbivores often reduce the abundance (biomass, cover) of dominant species [62, 63]. On the other hand, under high-nutrient availablity, the nutrient pulse and the herbivory treatment hardly affected the dominance of alien plants. One plausibility explanation for this finding is that the overall high-nutrient availability reduces, or even cancels, the nutrient-limitation shifts over time caused by nutrient fluctuations [20, 24, 64], resulting in very weak effects of the nutrient pulse on dominance changes in plant communities [65]. In addition to the weak evidence that herbivory mediated the effects of changes in nutrient availability and fluctuations on alien plant invasion, a recent case study also found that the parasitic plant *Cuscuta australis* could also regulate the effects of nutrient availablity and fluctuations on the invasion success of the alien plant *Bidens pilosa* [25](Gao et al. 2021). Therefore, if organisms from other trophic levels frequently mediate the effects of nutrient fluctuations on alien plant invasion, we recommend that studies testing the fluctuating resources hypothesis should more frequently consider the interactive effect of other trophic levels.

## Conclusions

The fluctuating resource hypothesis has become a key theory for explaining invasion success of alien plants. However, our study is, to the best of our knowledge, the first multi-species experiment that tested how another trophic level influences the effects of nutrient fluctuations on alien plant invasion. Partly in line with the fluctuating resource hypothesis, we found tentative evidence that nutrient variability promotes alien plant invasion only under overall low-nutrient conditions, and only in the absence of herbivores. Therefore, other trophic levels, such as herbivores in our study, might mediate the interactive effect of nutrient enrichment on alien plant invasion into resident communities.

## Acknowledgements

We thank Xue Zhang, Mingxin Pan, Chunling Chang, Huifei Jin, Lichao Wang, Lingxi Wang, Liping Shan, Han Yu for set-up of the experiment and plant harvest, and we thank Mingxin Pan for weighing the biomass. This work was supported by funding from the Chinese Academy of Sciences (Y9B7041001).

## Author contributions

Y Liu conceived the idea and designed the experiment. Y Li and Y Gao performed the experiment. Y Li and Y Liu analyzed the data. Y Li and Y Liu wrote the first draft of the manuscript, with further inputs from Y Gao and M van Kleunen.

## Data availability

All data and code available from the Dryad Digital Repository https://doi.org/10.5061/dryad.fj6q573vn (Li et al., 2021).

## Supporting information

### Methods S1

#### Recipe for 400% strength Hoagland’s Complete Nutrient Solution

The preparation of Hoagland nutrient solution essentially followed recommendations by Hoagland and Arnon (1950) and Liu et. al (2017), with the exception of the form in which iron was added (see below). We prepared the stock solutions 1-6 and micronutrient-stock solution given below, and used the amounts indicated to prepare 1 liter of nutrient solution:

1. 4 mL of 1.00 M/L KH_2_PO_4_
2. 20 mL of 1.00 M/L KNO_3_
3. 20 mL of 1.00 M/L Ca(NO_3_)_2_
4. 8 mL of 1.00 M/L MgSO_4_
5. 4 ml of micronutrient stock solution (see recipe below)
6. 10 ml of 1000 mg/liter iron from iron chelate (Fe-EDTA)

**\* Micronutrient-stock solution per liter:**

2.86 g H_3_BO_3_
1.81 g MnCl_2_·4H_2_O
0.22 g ZnSO_4_·7H_2_O
0.08 g CuSO_4_·5H_2_O
0.018 g H_2_MoO_4_ (Assaying 94% MoO_3_)

* Hoagland’s recipe called for 1 ml of 0.5% iron tartrate stock per liter of nutrient solution but we used iron chelate instead.

Reference: Hoagland, D.R. & Arnon, D.I. 1950. The water-culture method of growing plants without soil. California Agricultural Experiment Station, Circular 347.

**Table S1.** Detailed information on the eight alien target species and 15 native competitor species used in this study

| Species | Family | Status | Life history | Seed sources | Sowing date |
| --- | --- | --- | --- | --- | --- |
| *Bidens frondosa* L. | Compositae | Alien | Annual | Zhejiang | 26.06.2020 |
| *Bidens pilosa* L. | Compositae | Alien | Annual | Zhejiang | 26.06.2020 |
| *Solidago canadensis* L. | Compositae | Alien | Perennial | Zhejiang | 26.06.2020 |
| *Xanthium strumarium* L. | Compositae | Alien | Annual | Nei Mongol | 26.06.2020 |
| *Abutilon theophrasti* Medik. | Malvaceae | Alien | Annual | Jilin | 26.06.2020 |
| *Hibiscus trionum* L. | Malvaceae | Alien | Annual | Jilin | 05.07.2020 |
| *Lolium perenne* L. | Poaceae | Alien | Perennial | Greenwood flower seed company | 26.06.2020 |
| *Paspalum notatum* Flügge | Poaceae | Alien | Perennial | Zhejiang | 15.05.2020 |
| *Catolobus pendulus* (L.) Al-Shehbaz<sup>5</sup> | Brassicaceae | Native | Biennial | Nei Mongol | 26.06.2020 |
| *Cynanchum chinense* R. Br.<sup>2</sup> | Apocynaceae | Native | Annual | Nei Mongol | 26.06.2020 |
| *Lappula myosotis* V. Wolf<sup>1</sup> | Boraginaceae | Native | Annual or<br>Biennial | Nei Mongol | 26.06.2020 |
| *Artemisia rubripes* Nakai<sup>4</sup> | Compositae | Native | Perennial | Nei Mongol | 26.06.2020 |
| *Achillea asiatica* Serg.<sup>2</sup> | Compositae | Native | Perennial | Nei Mongol | 26.06.2020 |
| *Convolvulus arvensis* L.<sup>3</sup> | Convolvulaceae | Native | Perennial | Jilin | 26.06.2020 |
| *Gypsophila licentiana* Hand.-Mazz.<sup>4</sup> | Caryophyllaceae | Native | Perennial | Nei Mongol | 26.06.2020 |
| *Euphorbia humifusa* Willd.<sup>5</sup> | Euphorbiaceae | Native | Annual | Nei Mongol | 26.06.2020 |
| *Chloris virgata* Sw.<sup>2</sup> | Poaceae | Native | Annual | Nei Mongol | 26.06.2020 |
| *Cleistogenes squarrosa* (Trin.) Keng<sup>5</sup> | Poaceae | Native | Perennial | Jilin | 26.06.2020 |
| *Digitaria sanguinalis* (L.) Scop.<sup>1</sup> | Poaceae | Native | Annual | Jilin | 26.06.2020 |
| *Echinochloa crus-galli* (L.) P.Beauv.<sup>4</sup> | Poaceae | Native | Annual | Jilin | 26.06.2020 |
| *Setaria pumila* (Poir.) Roem. & Schult.<sup>3</sup> | Poaceae | Native | Annual | Jilin | 27.06.2020 |
| *Portulaca oleracea* L.<sup>3</sup> | Portulacaceae | Native | Annual | Nei Mongol | 26.06.2020 |
| *Rumex crispus* L.<sup>1</sup> | Polygonaceae | Native | Perennial | Nei Mongol | 26.06.2020 |

<sup>1,2,3,4,5</sup> Allocation of native species to the five native communities; each group included three different native species

**Table S2.** Output of the models residual standard deviations sigma for individual alien species. Shown are the model estimates and standard errors as well as the lower (L) and upper (U) values of the 95% and 90% credible intervals (CI).

|  |  | Estimate | SE | L95% CI | U95% CI | L90% CI | U90% CI |
| --- | --- | --- | --- | --- | --- | --- | --- |
| **Biomass production of alien target plants** | *Abutilon theophrasti* Medik. | -0.785* | 0.138 | -1.033 | -0.499 | -1.001 | -0.547 |
|  | *Bidens frondosa* L. | 1.074* | 0.127 | 0.839 | 1.341 | 0.874 | 1.290 |
|  | *Bidens pilosa* L. | 0.327* | 0.126 | 0.092 | 0.590 | 0.128 | 0.544 |
|  | *Hibiscus trionum* L. | -0.730* | 0.175 | -1.072 | -0.388 | -1.018 | -0.443 |
|  | *Lolium perenne* L. | 0.667* | 0.138 | 0.413 | 0.953 | 0.452 | 0.907 |
|  | *Paspalum notatum* Flügge | 0.444* | 0.123 | 0.217 | 0.700 | 0.249 | 0.659 |
|  | *Solidago canadensis* L. | -0.488* | 0.125 | -0.718 | -0.232 | -0.688 | -0.277 |
|  | *Xanthium strumarium* L. | 0.681* | 0.127 | 0.445 | 0.944 | 0.480 | 0.900 |
| **Biomass production of native communities** | *Abutilon theophrasti* Medik. | 0.373* | 0.133 | 0.135 | 0.652 | 0.169 | 0.602 |
|  | *Bidens frondosa* L. | 0.591* | 0.129 | 0.354 | 0.862 | 0.388 | 0.810 |
|  | *Bidens pilosa* L. | 0.511* | 0.125 | 0.275 | 0.766 | 0.310 | 0.725 |
|  | *Hibiscus trionum* L. | 0.603* | 0.127 | 0.370 | 0.869 | 0.403 | 0.822 |
|  | *Lolium perenne* L. | 0.484* | 0.146 | 0.216 | 0.788 | 0.253 | 0.737 |
|  | *Paspalum notatum* Flügge | 0.960* | 0.122 | 0.731 | 1.209 | 0.768 | 1.168 |
|  | *Solidago canadensis* L. | 0.721* | 0.113 | 0.510 | 0.949 | 0.539 | 0.911 |
|  | *Xanthium strumarium* L. | 0.563* | 0.123 | 0.335 | 0.816 | 0.369 | 0.773 |
| **Biomass proportion of alien target plants** | *Abutilon theophrasti* Medik. | 2.471* | 0.134 | 2.219 | 2.749 | 2.258 | 2.700 |
|  | *Bidens frondosa* L. | 2.559* | 0.133 | 2.314 | 2.833 | 2.353 | 2.783 |
|  | *Bidens pilosa* L. | 2.206* | 0.128 | 1.971 | 2.470 | 2.004 | 2.426 |
|  | *Hibiscus trionum* L. | 2.624* | 0.141 | 2.355 | 2.912 | 2.398 | 2.860 |
|  | *Lolium perenne* L. | 2.962* | 0.144 | 2.698 | 3.262 | 2.737 | 3.211 |
|  | *Paspalum notatum* Flügge | 2.908* | 0.123 | 2.679 | 3.162 | 2.712 | 3.116 |
|  | *Solidago canadensis* L. | 2.416* | 0.125 | 2.176 | 2.669 | 2.218 | 2.626 |
|  | *Xanthium strumarium* L. | 2.596* | 0.127 | 2.355 | 2.858 | 2.394 | 2.811 |

**Figure S1.**
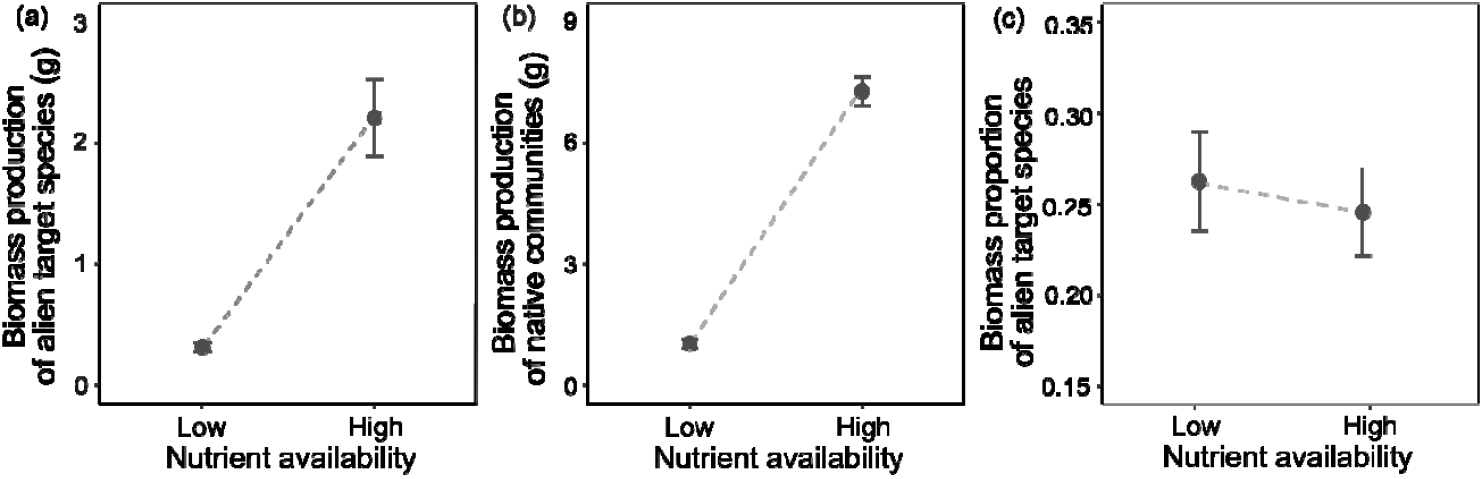
Mean values (±SE) of biomass production of the alien target species (a) and the native communities (b) and biomass proportion of the alien target species (c) under different nutrient availability conditions.

**Figure S2.**
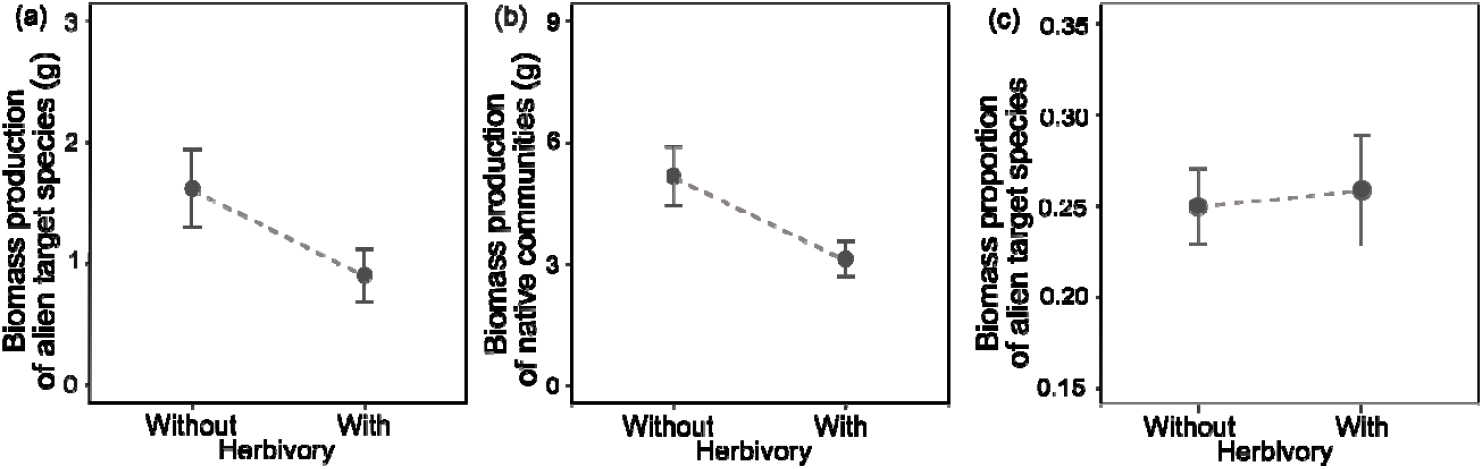
Mean values (±SE) of biomass production of the alien target species (a) and the native communities (b) and biomass proportion of the alien target species (c) growing with and without herbivory.

**Figure S3.**
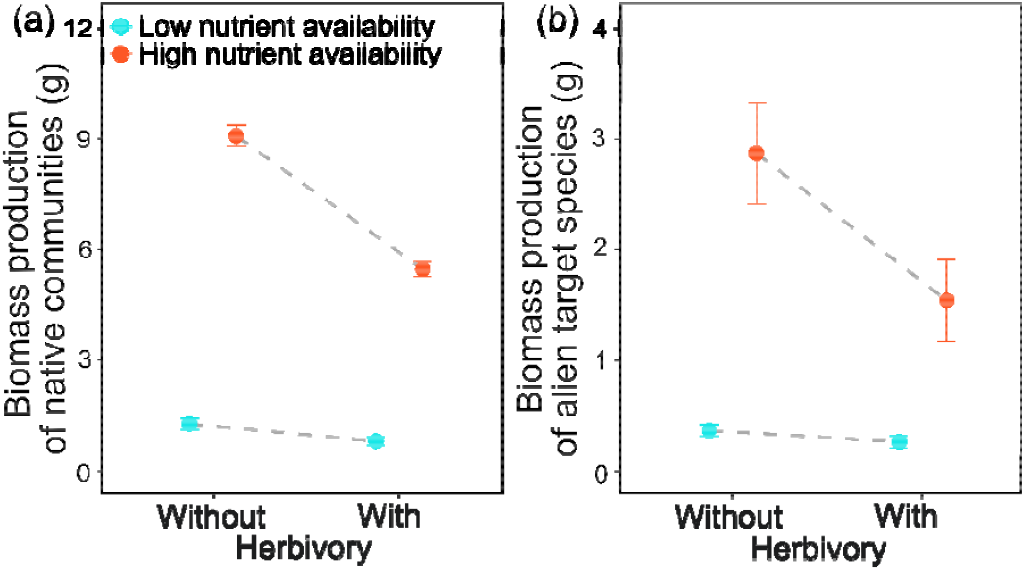
Mean values (±SE) of biomass production of native communities (a) and alien target species (b) under each combination of two nutrient availability (low *vs* high) and two herbivory (with *vs* without) treatments.

**Figure S4.**
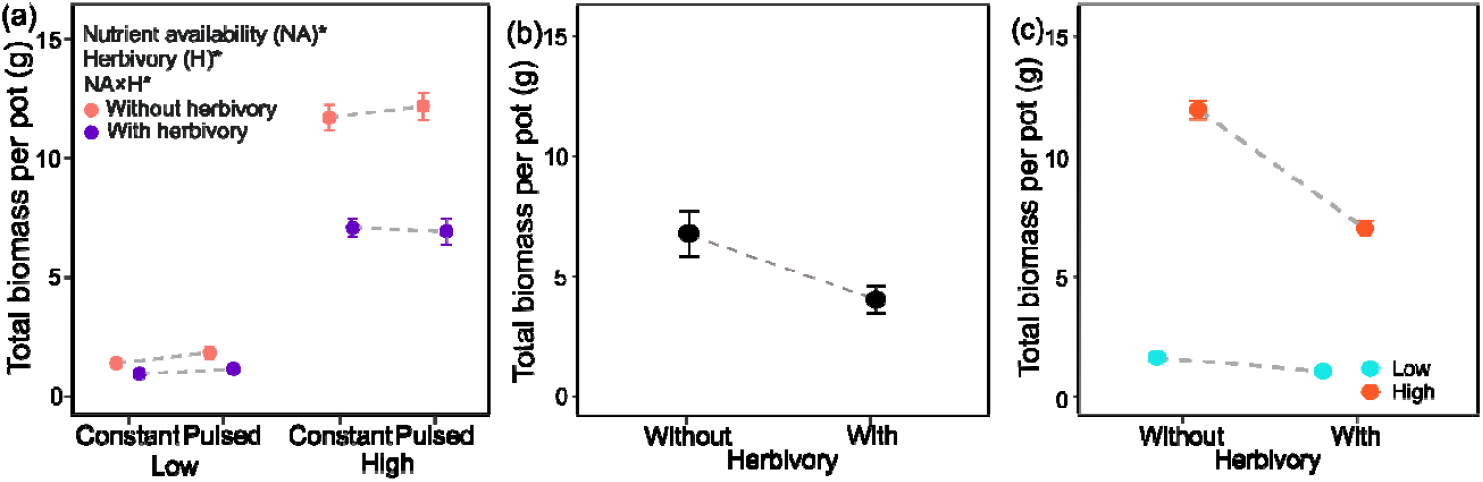
Mean values (±SE) of total biomass per pot under each combination of two nutrient availability (low *vs* high), two nutrient fluctuation (constant *vs* pulsed) and two herbivory (with *vs* without) treatments (a), with and without herbivory (b), and under each combination of two nutrient availability (low *vs* high) and two herbivory (with *vs* without) treatments (c). Parameters whose 95% credible intervals do not overlap with zero are indicted with asterisks (*).

